# Motor learning adapts to effort-dependent uncertainty

**DOI:** 10.64898/2026.09.10.750575

**Authors:** Polina E. Zhigulina, Lukas J. Volz, Tanya Bentley, Philip Zeyen, Frank Jessen, Marc Tittgemeyer, Lionel Rigoux

## Abstract

Adaptive behaviour depends on continuously updating motor control when action outcomes differ from expectations. Because action execution and sensory feedback are noisy, the value of an error depends on how reliably it reflects the consequences of an action. Most studies have manipulated the reliability of sensory feedback or action outcomes rather than uncertainty generated by the action itself. Here we show that motor adaptation is calibrated to effort-dependent uncertainty arising during action generation. Using a force-based motor adaptation paradigm, we demonstrate that greater effort progressively increased uncertainty in action execution and proprioceptive feedback, with corresponding changes in adaptation rate that closely matched normative predictions. Reward enhanced adaptation by improving proprioceptive precision rather than directly changing adaptation rate. Supporting this framework, patients with major depressive disorder exhibited elevated execution and proprioceptive uncertainty, impaired reward-dependent improvement in proprioceptive precision, and slower motor adaptation. These findings establish intrinsic uncertainty as a general determinant of adaptive behaviour, linking action generation, sensory evaluation, and motivation.

## Introduction

Neural transmission is both noisy and metabolically costly. Both the generation of voluntary actions and the estimation of their sensory consequences are therefore subject to constraints imposed by uncertainty and energetic demands ^1, 2^. How these intrinsic constraints shape the learning processes that continuously recalibrate behaviour remains incompletely understood ^3^.

Motor learning relies on internal predictions about the sensory consequences of actions and on continuous error-driven updating when outcomes deviate from expectations ^4, 5^. Within Bayesian and optimal control frameworks, adaptation rates are adjusted according to uncertainty in sensory feedback and motor execution ^6–8^. Experimental studies have demonstrated such uncertainty-dependent adaptation in response to altered sensory feedback and environmental perturbations ^9–11^, suggesting that motor adaptation is fundamentally governed by uncertainty-sensitive computations.

Whether the same principles govern uncertainty arising intrinsically during action generation remains largely unresolved. Work in animal models has established that motor variability is an intrinsic property of action generation and is relevant to motor learning ^12–14^. Human studies, meanwhile, have primarily examined uncertainty by imposing perturbations or altering sensory feedback, often conceptualising intrinsic motor variability as a relatively stable individual property ^7, 8^. Yet every action also generates its own inherent uncertainty. As motor commands are subject to signal-dependent noise, motor variability increases predictably with force production ^15^. Consequently, increasingly effortful actions are intrinsically more uncertain, providing a natural source of variability that differs fundamentally from experimentally imposed perturbations. Normative theories predict that adaptation should adjust to these fluctuations, with slower adaptation becoming optimal as intrinsic motor uncertainty increases ^7, 8^. Direct empirical evidence that motor adaptation is tuned to naturally occurring, effort-dependent uncertainty, however, is lacking to date.

Motor adaptation is also affected by rewards: positive and negative feedback have been reported to differentially influence aspects of motor learning and retention ^16, 17^, although the mechanisms are unclear. As reward reliably increases motor vigour ^18, 19^ and can modulate sensory precision ^20, 21^, motivation may influence adaptation by altering how the costs of action and the reliability of sensorimotor information are balanced. This possibility should be particularly relevant for effortful actions, for which both energetic costs and intrinsic uncertainty are increased. Yet whether rewards directly alter adaptation rates or instead change the reliability of the sensorimotor information used for learning remains also unresolved.

These considerations point to several computationally distinct routes through which intrinsic uncertainty may influence adaptation ^1, 22^. Motor execution determines how reliably an intended motor command is translated into an action, sensory feedback determines how accurately movement errors can be evaluated, and motivational processes influence both the willingness to invest effort and the precision of sensorimotor representations ^18, 23, 24^. Because these processes interact, existing paradigms have struggled to dissociate their respective contributions to adaptive motor behaviour. We therefore developed a force-based motor adaptation paradigm that systematically manipulates effort demands while independently varying reward and sensory feedback, allowing execution uncertainty, proprioceptive uncertainty and adaptation dynamics to be quantified within a common computational framework.

We applied this framework in healthy participants and patients with major depressive disorder, a condition characterised by psychomotor retardation and impaired motor performance ^25, 26^. Although depression is frequently associated with altered motivation and reward processing ^27, 28^, its motor symptoms are unlikely to arise from motivational deficits alone and may instead reflect alterations in the mechanisms governing action execution and adaptation ^29, 30^. Depression therefore provides a clinically relevant model for examining how changes in intrinsic motor uncertainty influence adaptive behaviour.

Using behavioural analyses and computational modelling, we separately quantified motivational biases, execution uncertainty, proprioceptive uncertainty and adaptation dynamics. We tested whether effort-dependent uncertainty arising during action generation and sensory evaluation governs motor adaptation. Specifically, we show that increasing effort increases sensorimotor uncertainty, to which adaptation rates adjust in accordance with normative predictions. Depression was associated with elevated motor uncertainty and correspondingly slower adaptation, indicating that apparent learning deficits largely arose from altered uncertainty rather than impaired adaptation mechanisms. Finally, reward enhanced adaptation under proprioceptive guidance by improving proprioceptive precision, an effect that was impaired in depression. Together, these findings identify effort-dependent uncertainty as the computational quantity linking motivation, motor control and adaptive learning.

## Results

### Increasing effort increases motor uncertainty

Motor adaptation is commonly studied by perturbing the external environment—for example, through altered visual feedback or visuomotor rotations. Conversely, we asked whether motor adaptation changes as increasing effort makes actions intrinsically more uncertain. To address this question, we developed a novel isometric force task in which participants learned to generate precise force pulses to move a virtual marble towards a target (Fig. 1a). Our force-based paradigm builds on previous work linking force production to incentive motivation and motor vigour ^18, 19^. However, we here exploit the force-dependent increase in motor variability to parametrically manipulate intrinsic uncertainty; the expected incentive effect on vigour was also replicated in our data (Extended Data Fig. 1). Because motor variability scales with force production, increasing the required force naturally increased motor uncertainty while leaving the external environment unchanged.

**Figure 1.**
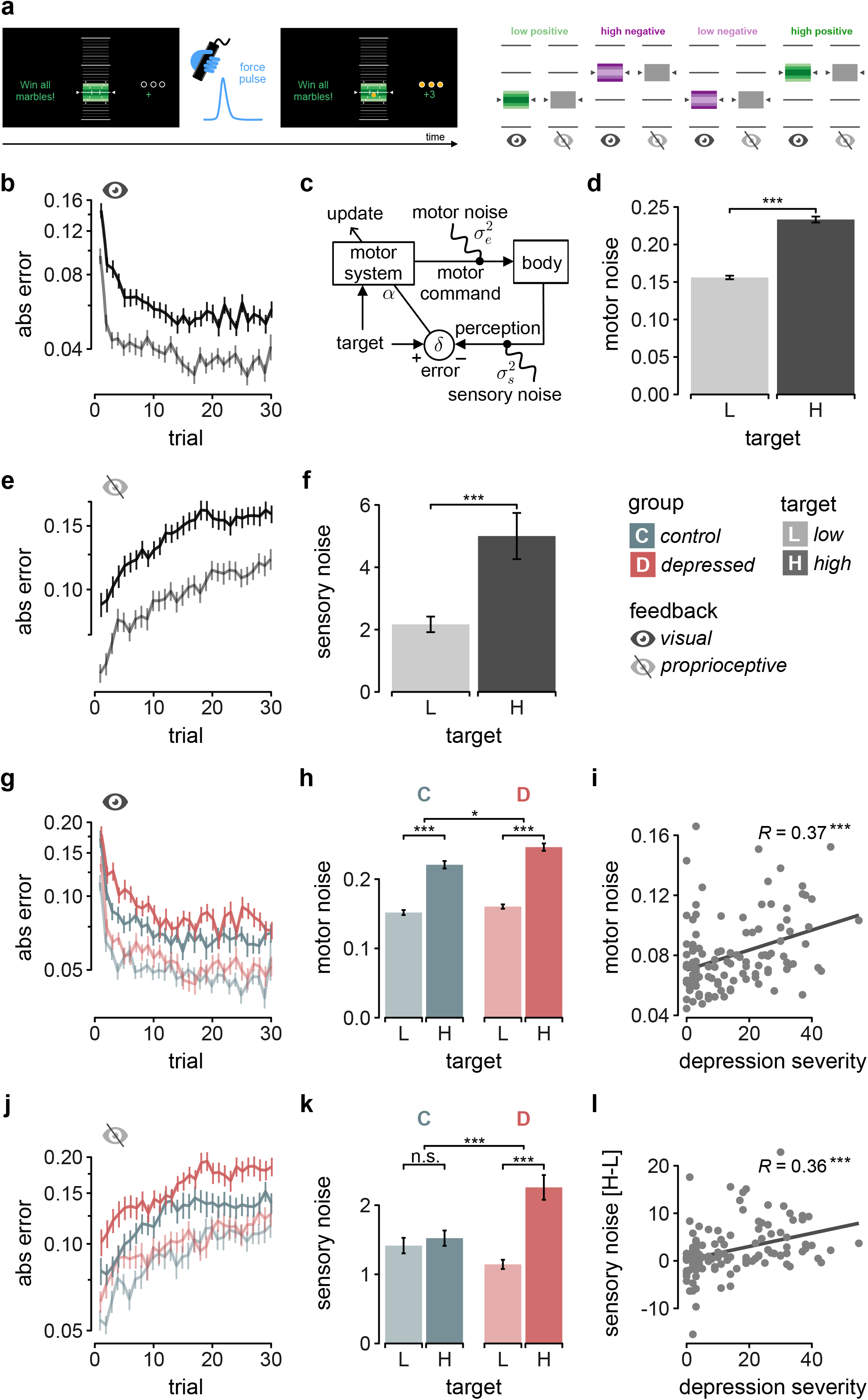
Motor execution and sensory uncertainty increase with effort and are elevated in depression. a, Experimental design. Participants produced brief force pulses to move a marble towards a target at low or high force requirements under visual or proprioceptive feedback, with positive or negative reinforcement. b, Learning curves for absolute aiming error under visual feedback for low- and high-target conditions. c, State-space model used to estimate motor execution noise, sensory noise and adaptation rate. d, Motor execution noise estimated from the model for low- and high-target conditions. e, Aiming-error drift following withdrawal of visual feedback. f, Sensory noise estimated from the model under proprioceptive feedback. g,h, Learning curves and motor execution noise under visual feedback for controls and patients in low and high effort conditions. i, Relationship between motor execution noise and depression severity. j,k, Aiming-error drift and sensory noise under proprioceptive feedback for controls and patients in low and high effort conditions. l, Relationship between the change in sensory noise across target conditions and depression severity. Low-target (L) and high-target (H) conditions are shown in lighter and darker shades, respectively; healthy controls (C) and depressed patients (D) are shown in blue and red. Data are means ± SEM. Statistical tests and exact p-values are reported in the main text. Significance levels: p* < .05, p*** < .001.

Participants (N = 121; 61 healthy controls, 60 with major depressive disorder; groups were matched for age, p < .956, and sex, p > .416) performed the task at two force levels corresponding to 20% and 40% of their maximal voluntary force. Participants alternated between blocks with visual feedback and blocks without visual feedback during which participants relied on proprioception alone. Patients exhibited marked psychomotor retardation (SRRS: t = 15.2, p < .001, Table 1), allowing us to test whether the same computational principles extended to pathological motor behaviour.

**Table 1.** Summary of the demographics and clinical variables.

|  | **C (N=61)** | **D (N=60)** | **Total (N=121)** | **p-value** |
| --- | --- | --- | --- | --- |
| **age** |  |  |  | 0.956 |
| Mean (SD) | 30.738 (7.380) | 30.833 (11.341) | 30.785 (9.512) |  |
| Range | 20.000 - 54.000 | 18.000 - 65.000 | 18.000 - 65.000 |  |
| **sex** |  |  |  | 0.416 |
| F | 26 (42.6%) | 30 (50.0%) | 56 (46.3%) |  |
| M | 35 (57.4%) | 30 (50.0%) | 65 (53.7%) |  |
| **MADRS** |  |  |  | < 0.001 |
| N-Miss | 14 | 9 | 23 |  |
| Mean (SD) | 1.021 (1.452) | 16.882 (8.701) | 9.276 (10.171) |  |
| Range | 0.000 - 6.000 | 0.000 - 37.000 | 0.000 - 37.000 |  |
| **BDI-II** |  |  |  | < 0.001 |
| Mean (SD) | 3.557 (3.299) | 25.400 (10.704) | 14.388 (13.492) |  |
| Range | 0.000 - 12.000 | 0.000 - 55.000 | 0.000 - 55.000 |  |
| **DARS** |  |  |  | < 0.001 |
| Mean (SD) | 57.525 (6.310) | 44.217 (11.104) | 50.926 (11.188) |  |
| Range | 36.000 - 67.000 | 8.000 - 66.000 | 8.000 - 67.000 |  |
| **SRRS** |  |  |  | < 0.001 |
| Mean (SD) | 2.066 (1.999) | 12.600 (4.982) | 7.289 (6.494) |  |
| Range | 0.000 - 7.000 | 2.000 - 24.000 | 0.000 - 24.000 |  |
| **SRRS-motor** |  |  |  | < 0.001 |
| Mean (SD) | 1.180 (1.397) | 4.667 (2.735) | 2.909 (2.778) |  |
| Range | 0.000 - 5.000 | 0.000 - 11.000 | 0.000 - 11.000 |  |

Participants rapidly reduced aiming errors across trials, demonstrating robust motor adaptation (Fig. 1b). Adaptation, however, depended strongly on effort: errors remained larger for the high-force condition (Extended Data Fig. 2a-b) and declined more slowly across learning (target: F = 269.1, p < .001; trial × target: F = 17.5, p < .001). Thus, increasing effort affected both the magnitude of motor errors and how they were corrected over time.

To determine which components of motor uncertainty accounted for this effect, we decomposed behaviour into execution uncertainty, proprioceptive uncertainty and adaptation rate using a state-space model (Fig. 1c; Extended Data Fig. 2c; see Methods). Execution uncertainty and adaptation rate were estimated from the learning trajectories, whereas proprioceptive uncertainty was estimated from the subsequent phases without visual feedback. This allowed us to quantify these components within the same task. Execution uncertainty increased with force (t = 3.4, p < .001; Fig. 1d).

Sensory uncertainty scales with force. Without visual feedback, performance drifted progressively (F = 525.7, p < .001; Fig. 1e), with larger errors for the high target (t = 5.5, p < .001; Extended Data Fig. 2d-e) that also unfolded differently in time (trial × target: F = 125.3, p < .001). Because execution uncertainty was expected to produce variability rather than a progressive drift, the latter indicates increasing uncertainty in estimating movement errors as force increases ^24^. Consistent with this interpretation, the model estimated greater proprioceptive variance for the high target (t = 4.73, p < .001; Fig. 1f).

#### Depression amplifies sensorimotor uncertainty

Depression altered adaptation dynamics (trial × group: F = 4.8, p = .029; Fig. 1g; Extended Data Fig. 2f-l). The force-dependent increase in execution uncertainty was greater in patients than in healthy participants (group × target: F = 6.2, p = .014; Fig. 1h) and scaled with symptom severity (BDI-II; R = 0.37, p < .001; Fig. 1i; Supplementary Table 1). Although muscle mass predicted execution uncertainty (R = –0.24, p = .008; Extended Data Fig. 3a-b), its weak relationship with symptom severity (Extended Data Fig. 3b; Supplementary Table 1) suggests that differences in muscle mass do not account for this clinical association. Proprioceptive uncertainty showed a similar pattern: the progressive drift was greater for the higher target in patients (trial × target × group: F = 7.9, p = .005; Fig. 1j; Extended Data Fig. 2m-p), and the force-dependent increase in sensory variance was driven by the patient group (target × group: F = 18.5, p < .001; Fig. 1k). Importantly, proprioceptive variance also tracked clinical severity (BDI-II; R = 0.36, p < .001; Fig. 1l; Supplementary Table 1).

Together, these findings show that increasing effort raises both execution and proprioceptive uncertainty, with both sources of uncertainty amplified in depression. We next asked how motor adaptation is tuned to account for this uncertainty.

### Adaptation rates are tuned to motor uncertainty

A learner facing greater error uncertainty should correct less, not more: over-correcting noise-driven errors injects variability rather than reducing it ^7^. Previous work has shown that adaptation rates are adjusted to uncertainty introduced by external perturbations, such as perturbed or degraded visual feedback ^6, 31^. Whether the same principle extends to intrinsic uncertainty associated with increasingly effortful actions has remained unknown ^32^. We therefore asked whether adaptation rates decreased when intrinsic uncertainty increased.

Indeed, error correction was slower for the high target (t = 3.44, p < .001; Fig. 2a), and slower when visual feedback was removed (t = −13.3, p < .001; Fig. 2b). Within participants, the effort-driven change in execution precision (inverse variance; see Methods) was associated with the change in adaptation rate (R = 0.57, p < .001; Fig. 2c), and the same relationship was observed for sensory precision (R = 0.42, p < .001; Fig. 2d). Across participants, both execution precision (R = 0.36, p < .001; Fig. 2e) and sensory precision (R = 0.25, p = .006; Fig. 2f) predicted individual adaptation rates.

**Figure 2.**
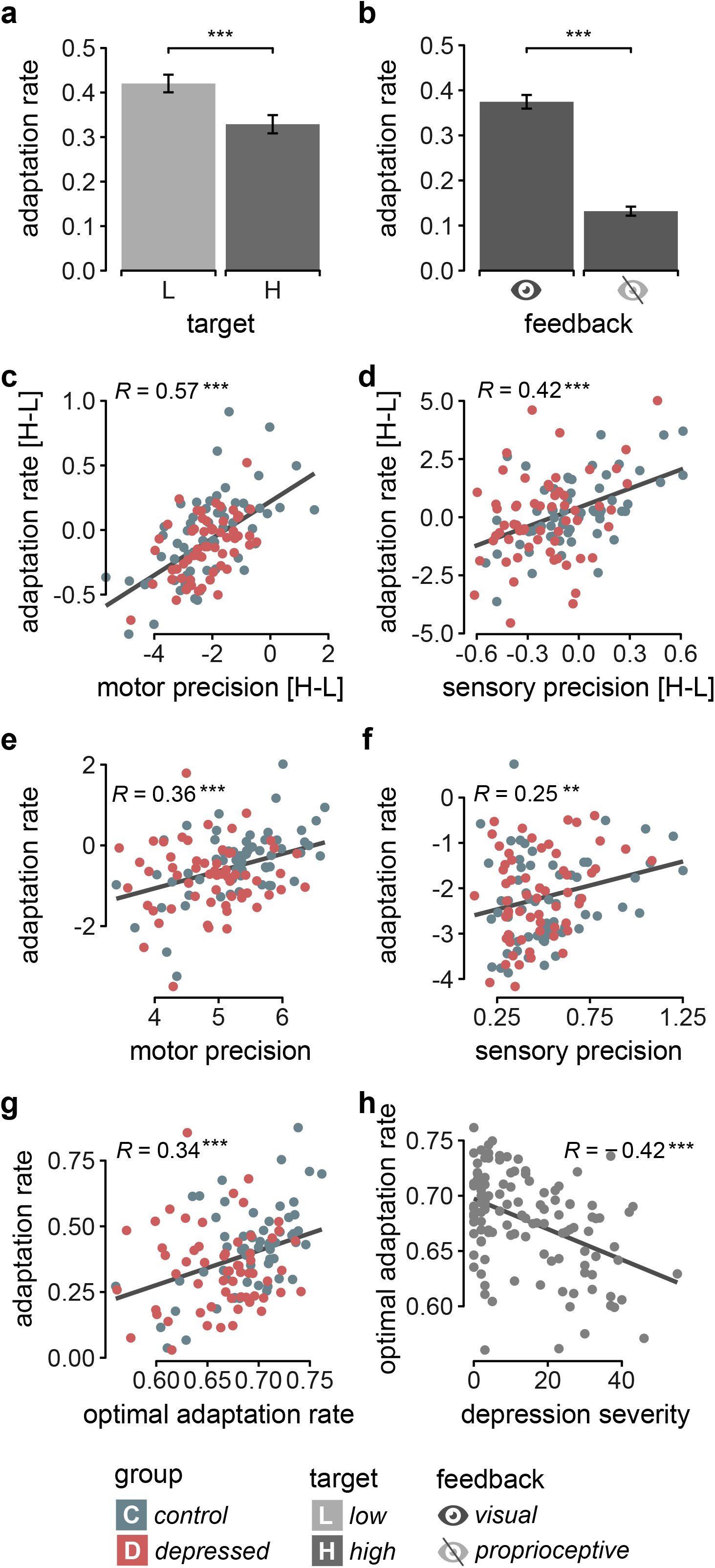
Adaptation rates depend on motor and sensory uncertainty. a, Adaptation rate estimated from the state-space model for low- and high-target conditions. b, Adaptation rate under visual and proprioceptive feedback. c,d, Relationship between the change in adaptation rate across target conditions and the corresponding change in motor and sensory precision, respectively. e,f, Relationship between adaptation rate and motor and sensory precision across participants. g, Relationship between observed and theoretically optimal adaptation rates. Optimal adaptation rate was derived from the model as the rate that minimises asymptotic error given the levels of motor and sensory uncertainty. h, Relationship between optimal adaptation rate and depression severity (BDI-II score). Motor precision is the inverse of the motor execution noise parameter estimated under visual feedback; sensory precision is the inverse of the sensory noise parameter estimated under proprioceptive feedback. Low-target (L) and high-target (H) conditions are shown in lighter and darker shades, respectively; healthy controls (C) and depressed patients (D) are shown in blue and red. Data are means ± SEM. Pearson’s correlations are shown with maximum-likelihood regression lines. Statistical tests and exact p-values are reported in the main text. Significance levels: p** < .01, p*** < .001.

Motor uncertainty explains slower adaptation in depression. Patients adapted more slowly than healthy controls (t = 2.7, p = .007; Extended Data Fig. 3d). To test whether elevated uncertainty accounted for this group difference, we used the estimated execution and sensory uncertainty to derive the optimal adaptation rate that minimises expected long-term error (see Methods). Participants closely followed these normative predictions (R = 0.34, p < .001; Fig. 2g). Moreover, the uncertainty-derived optimal adaptation rate was negatively associated with symptom severity (BDI-II: R = −0.42, p < .001; Fig. 2h; Supplementary Table 1), indicating that the elevated uncertainty in depression explains the observed slowing of adaptation. Thus, the slower adaptation observed in depression can be understood as an appropriately calibrated response to greater sensorimotor uncertainty.

#### Reward does not modulate adaptation rates

Reward is known to enhance aspects of motor performance and learning ^16, 21, 23^, raising the possibility that blunted reward sensitivity contributes to slower adaptation in depression. With visual feedback, however, neither overall performance (all p > .163; Fig. 3a) nor adaptation rates (t = – 0.55, p = .585; Fig. 3b) differed significantly between feedback conditions. The absence of a detectable effect on adaptation rates is not necessarily at odds with reports that incentives improve motor performance. Those effects may primarily act on online correction ^20^ rather than on updating the motor plan ^17^. Of note, our task used short ballistic pulses with no real-time visual guidance, providing little opportunity for such feedback-control mechanisms to operate.

**Figure 3.**
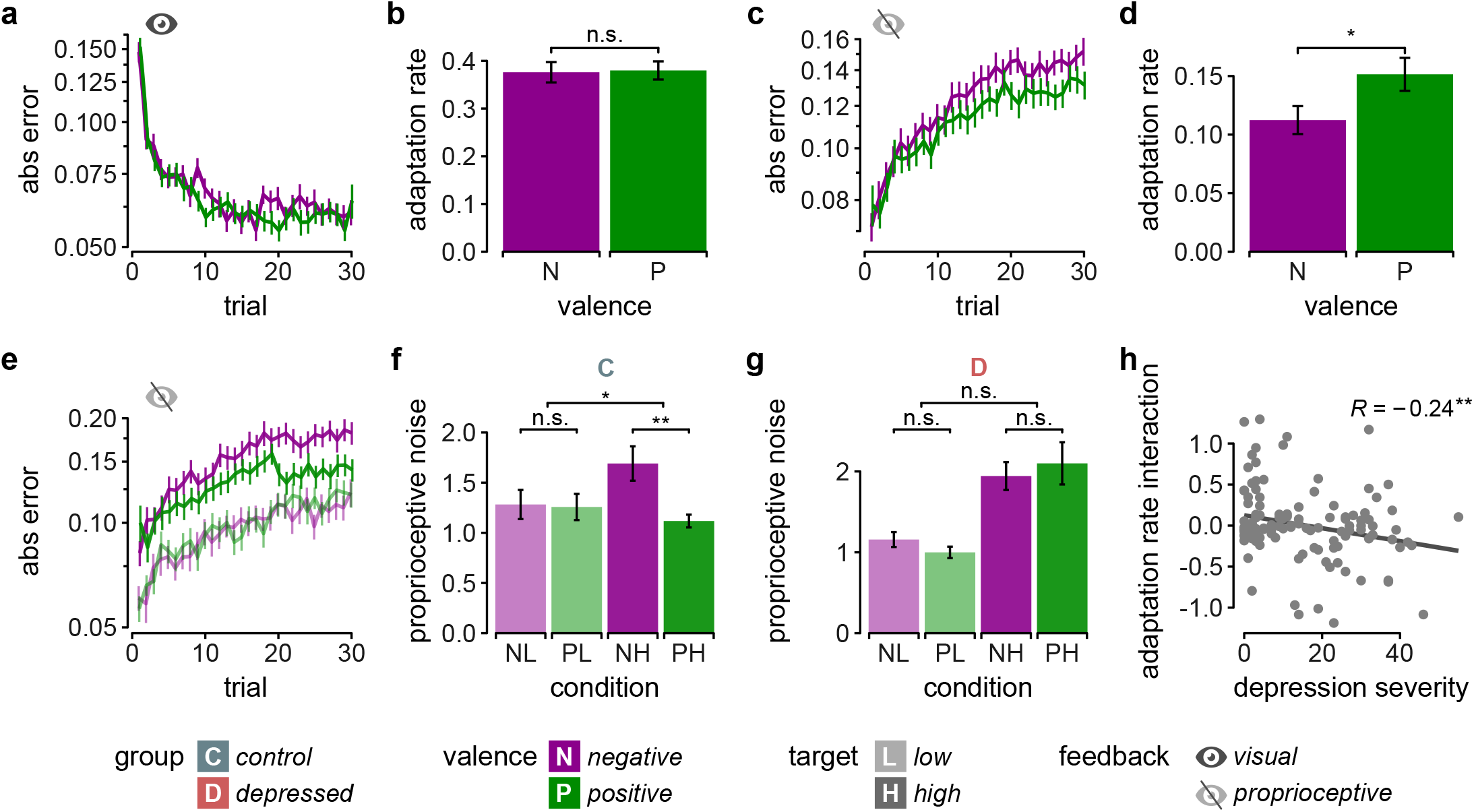
Reward improves proprioceptive precision during effortful motor adaptation. a, Aiming-error trajectories during adaptation under visual feedback for positive- and negative-valence conditions. b, Adaptation rate under visual feedback for positive- and negative-valence conditions. c, Aiming-error trajectories without visual feedback for positive- and negative-valence conditions. d, Adaptation rate without visual feedback for positive- and negative-valence conditions. e, Aiming-error trajectories without visual feedback for positive- and negative-valence conditions, shown separately for low- and high-target conditions. f, Estimated sensory noise without visual feedback in healthy controls, shown for low- and high-target conditions and positive- and negative-valence conditions. g, Estimated sensory noise without visual feedback in patients with depression, shown for low- and high-target conditions and positive- and negative-valence conditions. h, Relationship between depression severity (BDI-II score) and the effort × valence modulation of adaptation rate. The effort × valence modulation was calculated as the difference between the positive- and negative-valence changes in adaptation rate from low to high target. Negative-valence (N) and positive-valence (P) conditions are shown in purple and green; low-target (L) and high-target (H) conditions are shown in lighter and darker shades, respectively; healthy controls (C) and patients with depression (D) are shown in blue and red. Data in a–g are means ± SEM across participants. Pearson’s correlation is shown with a maximum-likelihood regression line. Statistical tests and p-values are reported in the main text. Significance levels: p* < .05, p** < .01.

Taken together, adaptation rates were tuned to motor uncertainty, whereas we found no evidence that reward altered adaptation rates when visual feedback was available.

### Reward enhances adaptation by improving proprioceptive precision

Under proprioceptive guidance, by contrast, reward did shape adaptation: performance drifted less following positive compared to negative feedback (trial × valence: F = 10.3, p = .001; Fig. 3c), and adaptation rates were higher following positive feedback (t = 2.1, p = .036; Fig. 3d). We therefore asked whether this effect of reward on adaptation arises from improved precision of the sensory signal on which adaptation depends.

Because proprioceptive uncertainty increased with force (Fig. 1f), we examined whether reward effects likewise varied with effort. The behavioural benefit of positive feedback was greater for higher effort (trial × valence × target: F = 9.74, p = .002; Fig. 3e). The model indicated that this effect arose from the sensory component: in controls, reward reduced the force-dependent increase in proprioceptive variance (target × valence: F = 4.4, p = .038; Fig. 3f). Thus, reward enhanced adaptation under proprioceptive uncertainty by improving the precision of the sensory signal on which adaptation depends.

### The reward-dependent improvement in proprioceptive precision is impaired in depression

Reward insensitivity is a hallmark of depressive disorder ^28^, and meta-analyses suggest that deficits are more pronounced for reward-driven learning and bias than for reward-driven vigour ^33, 34^. We therefore expected patients to show a reduced ability to modulate sensory uncertainty in response to reward, rather than a general reduction in reward-driven performance. The valence effect on performance indeed differed between groups (Extended Data Fig. 4a). In particular, the increase in sensory precision with positive feedback observed in healthy participants was absent in patients (target × valence: F = 0.9, p = .34; Fig. 3g; Extended Data Fig. 4b-e; group × target × valence: F = 4.2, p = .041; Extended Data Fig. 4f-h); the magnitude of the reward-related modulation of adaptation declined with symptom severity (BDI-II; R = – 0.30, p = .002; Fig. 3h; Table S1). Thus, patients not only experience greater motor uncertainty but also demonstrated a reduced capacity to improve proprioceptive precision when uncertainty was high. This combination provides a mechanistic account of slower adaptation in depression: elevated uncertainty appropriately reduces adaptation rates, while impaired reward-dependent sensory precision limits the ability to compensate for that uncertainty.

Together, these findings show that motor adaptation is tuned to uncertainty arising throughout the sensorimotor system, from action generation to action evaluation.

## Discussion

Motor adaptation has traditionally been studied under conditions in which uncertainty is imposed by the external environment ^35–37^. Here, we show that adaptation is calibrated to uncertainty arising intrinsically during action generation. Because motor variability increases with force production, increasingly effortful actions become progressively less reliable, resulting in decreased adaptation rates across effort levels, feedback conditions, and participants.

The same computational principle also extends to pathological motor behaviour. Patients with major depressive disorder exhibited elevated execution and proprioceptive uncertainty and adapted more slowly than healthy controls. Yet, across participants, adaptation rates closely followed the optimal rates predicted from their execution and proprioceptive uncertainty, and greater uncertainty in depression was sufficient to predict the observed difference in adaptation rates. Thus, in depression, impaired motor adaptation may largely reflect elevated motor uncertainty rather than a fundamental deficit in the adaptation process itself, while impaired reward-dependent improvement in proprioceptive precision may further limit compensation for that uncertainty.

These findings extend normative theories of motor adaptation to a source of uncertainty that has received comparatively little experimental attention. Bayesian accounts predict that adaptation should weight errors less strongly when motor output is unreliable, and this prediction has been supported for uncertainty introduced experimentally through altered sensory feedback or visuomotor perturbations ^7, 8, 31^. Whether the same principle applies to uncertainty arising intrinsically during action generation has remained unresolved. Unlike externally imposed perturbations, intrinsic uncertainty changes with the action itself and therefore cannot be varied independently of movement. By exploiting the signal-dependent increase in motor variability with force production ^15, 38, 39^, we found that adaptation rates tracked participants’ motor uncertainty, consistent with adaptation calibrating to the reliability of action outcomes. This distinction complements a large literature showing that centrally generated neural variability can facilitate motor learning by promoting exploration and increasing the flexibility of motor plans ^13, 23, 40, 41^. Notably, this planning variability serves a different computational role from the execution and proprioceptive uncertainty examined here: rather than providing exploratory variation, these sources of uncertainty limit the reliability of the error signal available for adaptation and may therefore lead to more conservative error correction.

Our findings also suggest a novel explanation for how reward influences motor adaptation, an issue that remains unresolved. Reward did not affect adaptation rates when visual feedback was available, but it enhanced adaptation under purely proprioceptive guidance, where sensory uncertainty was greater. This conditional effect may reconcile our findings with previous reports of beneficial effects of incentives on motor performance ^20, 21^. By using brief ballistic force pulses with feedback only after movement completion, our paradigm isolated between-trial adaptation from online motor correction, a distinction recently emphasised experimentally ^17^. This distinction may also help explain previous reports that positive feedback preferentially improves motor retention rather than adaptation ^16^. Enhanced proprioceptive precision could stabilise motor memories without requiring a change in the adaptation rate itself.

The mechanisms underlying this reward-dependent improvement in proprioceptive precision remain unclear. One possible explanation is that incentive motivation reduces neural variability by increasing investment in computational resources, as proposed by the cost-of-precision framework ^18, 21, 42^. Reward may thereby justify the energetic cost of generating more precise sensorimotor representations, improving the quality of the error signal available for adaptation.

Alternatively, reward may enhance proprioceptive precision through peripheral mechanisms. Co-contraction of antagonist muscles increases limb stiffness and proprioceptive acuity, reducing movement variability at a metabolic cost ^43–45^, and is recruited preferentially during the early stages of adaptation before internal models become fully established ^46^. Although these mechanisms operate at different levels of the motor system, both predict a selective precision benefit when uncertainty is sufficiently high to justify the additional energetic investment, consistent with our findings. Both mechanisms imply that motor precision is not fixed by the effector but can be actively regulated. This view is consistent with animal studies showing that movement variability arises in part from central rather than purely peripheral sources ^12^.

This may also explain why reward-dependent improvements in proprioceptive precision were absent in depression. If the energetic or computational cost of increasing precision is elevated, the same motivational signal may be less effective in recruiting the processes that improve sensory precision. The observation that dopaminergic function modulates the cost of precision in Parkinson’s ^21^ disease provides a potential link between motivational control of sensory precision and the dopaminergic dysfunction associated with depression ^28^.

These findings offer a novel interpretation of psychomotor dysfunction in depression. Psychomotor retardation is among the most robust observable features of the disorder ^26, 30, 47^, and slower adaptation can readily be interpreted as a deficit in motor learning. However, our findings suggest that the adaptation mechanism remains appropriately calibrated to the underlying uncertainty. Adaptation slowing tracked psychomotor retardation more closely than overall symptom severity and was accounted for by elevated execution and proprioceptive uncertainty under a common adaptation policy. At the same time, patients lacked the reward-dependent increase in proprioceptive precision observed in healthy participants, and this impairment increased with symptom severity. Thus, patients face both greater uncertainty and a reduced capacity to compensate for it: elevated uncertainty appropriately slows adaptation, while impaired reward-dependent sensory precision limits the ability to reduce that uncertainty. Viewed in this way, psychomotor retardation is less consistent with a primary deficit of motor adaptation than with the computational consequences of elevated motor uncertainty combined with impaired reward-dependent sensory precision. This distinction may have implications for interventions aimed at improving motor function in depression.

Several limitations qualify the interpretation of our findings. The clinical test of this framework in major depressive disorder was cross-sectional, and although adaptation slowing tracked psychomotor retardation more closely than overall symptom severity, we cannot determine whether elevated motor uncertainty precedes the motor syndrome or emerges as its consequence. Medication effects likewise cannot be separated from illness-related changes in the present cohort.

Our computational model assumes a constant adaptation rate and therefore does not distinguish between the interacting fast processes that can contribute to short-term motor adaptation ^48^. This abstraction was sufficient to capture the systematic relationship between motor uncertainty and adaptation rate, but does not address whether un-certainty differentially affects these processes. Furthermore, planning uncertainty comprises multiple computational components ^13, 49^. We minimised these influences through a highly predictable blocked design and therefore did not quantify them independently.

More broadly, our findings identify effort-dependent uncertainty as a computational link between motor execution, sensory precision, and adaptive behaviour. They show that adaptation rate is calibrated to the uncertainty under which behaviour is generated and evaluated. Intrinsic uncertainty therefore provides a general mechanism through which physiological state can shape motor learning, with the same principle potentially extending to conditions such as Parkinson’s disease, where dopaminergic dysfunction alters motor variability and the cost of precision ^50^, and healthy ageing, where increasing neuromotor noise can be accompanied by preserved or even exaggerated behavioural caution ^51, 52^. Across these conditions, our framework makes a clear prediction: slower adaptation may reflect not impaired learning, but the appropriate computational response of an adaptive system operating under greater intrinsic uncertainty.

## Methods

### Participants

A cohort of 60 individuals (26 women, 30.7±7.3 years) with unipolar major depressive disorder as defined in the ICD-10 was recruited at the Department of Psychiatry and Psychotherapy of the University of Cologne. In addition, 61 healthy controls (30 women, 30.8±11.3 years) were recruited from a pre-existing database from the Max Planck Institute for Metabolism Research.

We only included participants aged 18 years or older. We excluded participants with moderate to high suicidality risks Columbia Suicide Severity Rating Scale ^53^, signs of dementia from a brief cognitive Dementia Detection Test (score < 9; DemTect ^54^), known psychiatric or neurological comorbidities, as well as autoimmune diseases. We also excluded individuals taking benzodiazepines, antipsychotic, or anti-inflammatory medications, as well as those receiving electroconvulsive therapy. Treatment with antidepressants was accepted. Participants consuming nicotine (>10 cigarettes per day), or who were pregnant or breastfeeding, were also excluded.

All participants provided written informed consent before inclusion and underwent the same experimental procedure at the Max Planck Institute. The Ethical Committee of the Medical Faculty of the University of Cologne approved the study protocol (No. 20-1523_1).

### Clinical Assessments

A trained psychiatrist screened all participants for major depressive disorders according to the ICD-10 and using standardised scales. General depression severity was measured using the Montgomery–Åsberg Depression Rating Scale, MADRS ^55^, a gold standard for assessing depression in clinical trials ^56^. Participants also self-rated their depression severity using the Beck‘s Depression Inventory (BDI-II ^57^). Psychomotor retardation was assessed using the clinician-rated Salpetière Retardation Rating Scale (SRRS ^58^); items 1-5, specific to motor domains, and item 15, a global evaluation of psychomotor retardation, were pooled to derive a motor retardation score excluding cognitive domains. Anhedo-nia was measured with the self-rated Dimensional Anhedonia Rating Scale, DARS ^59^designed to assess desire, motivation, and consummatory pleasure across hedonic domains. Here, a high score reflects more intense reward sensitivity, i.e., low anhedonia. For all other scales, larger scores indicate more severe symptoms.

### Motor Learning Task

To assess motor learning behaviour, we implemented a new computerised task requiring participants to squeeze an isometric digital dynamometer (HD-BTA, Vernier™, Beaverton, OR). The force-based task was adapted from established paradigms used to investigate the influence of incentive motivation on motor vigour ^18, 19^. In the present study, we parametrically varied intrinsic uncertainty by manipulating force amplitude during action generation.

After reading the task instructions, participants first completed 3 maximal voluntary force (MVF) measurements to calibrate task difficulty to their individual force abilities. During each 3s attempt, real-time force output and the participant’s previous best record were displayed on-screen as numerical values (in Newton).

The task was framed as a “marble game” (see Fig. 1a) in which the participant had to throw “marbles” up a graduated scale on the screen with the goal of hitting a coloured target to win (or avoid losing) more marbles. On each trial, a short (< 700 ms) force pulse applied to the dynamometer animated a virtual marble moving up from the bottom of the screen and stopping at a position determined by the amplitude of the force pulse. Participants were instructed to earn as many marbles as possible and were informed that their global performance would determine how much money they would receive at the end of the testing day.

The target, indicated by white flanking arrows, was located at either 1/3 (low) or 2/3 (high) of the scale, representing respectively 20% and 40% of the participant’s MVF. Shaded areas around the target (±4.5%, ±3%, and ±1.5% of the MVF) delineated “hit zones” defining the outcome of the trial: When the target was green (“win” condition), the participant could respectively earn 1, 2, or 3 marbles by landing on the respective hit zones (and nothing if the marble landed outside of the largest one); when the target was purple (“loss” condition), hitting the central zone was the only way to avoid losing marbles, each step further withdrawing one additional marble from the total score, up to 3 marbles when the marble completely missed the hit zones. At the end of each trial, the hit zone on which the marble landed flickered, and the number of won/lost marbles was displayed on the side of the scale to provide additional feedback to the participant.

The task was organised in 4 blocs corresponding to the 4 possible combinations of target level (high vs low) and outcome valence (win vs loss). The order of the blocs was counterbalanced within each group. Each block consisted of a sequence of 30 identical trials, allowing participants to gradually improve their performance (learning phase under visual feedback). To assess participants’ ability to maintain the learned motor policy without visual outcome feedback, they performed 30 additional retention trials (i.e. without visual and only proprioceptive feedback) in which the marble was displayed at the target irrespective of the participant’s actual force output. Participants were explicitly informed that the displayed marble position was uninformative about their performance. The outcome was replaced by question marks, and the grey hit zone and semitransparent marble indicated that visual feedback about movement accuracy was unavailable. The task thus consisted of 240 trials (4 blocs of 30+30 trials). Trials started with the presentation of the target, and participants self-paced their response. Visual feedback (marble animation, target and outcome flickering) lasted a total of 4.6 s, and a jittered inter-trial interval (1.3 – 2.0 s, uniformly distributed) mitigated anticipatory effects.

Trials with faulty execution (reaction time < 50 ms, “too early”, or force pulse duration > 700 ms, “too slow”) were aborted and repeated after a warning message was shown; only valid trials contributed to the predefined trial count. To learn how to produce adequate force pulses, participants started with a short training session that lasted until they performed seven consecutive correct trials. They then completed a single trial of each condition—including with blinded feedback—as examples (order randomised) before starting the main trial sequence.

The task took approximately 35 minutes to be completed. Final total gain (5 € + 0.05 € per earned marble, minimum 3 €) was displayed on-screen at the very end.

The task was implemented in MATLAB 2019a using the Psychtoolbox v3.0.14.

### Statistical analysis

Signed aiming errors were computed as the difference between the marble endpoint and the target position on the scale (both between 0 and 1). Absolute aiming error was obtained from the absolute value of this difference. To aggregate behaviour across target levels, we computed the normalised error as the absolute aiming error divided by the square root of target position, effectively standardising performances to a comparable level.

Statistical analyses were implemented in R Statistical Software (v4.3.0) ^60^. To account for repeated measures within participants (trials and conditions), behavioural measures were fitted using linear mixed-effects models (including random intercepts for each participant), which were then analysed with an ANOVA procedure as implemented in R’s lmerTest package (v3.1.3) ^61^. Follow-up statistical tests for group and condition effects were performed using unpaired and paired two-sample Student’s t tests, and corrected for multiple comparisons using the Bonferroni method. Pearson’s correlations were used to identify relationships between clinical, behavioural, and computational metrics. Significance threshold was set to p < .05.

### Motor learning model

We modelled the evolution of force pulse amplitude (i.e., the marble endpoint) across trials using a state-space model ^62, 63^, in which motor noise arising during planning is distinguished from noise arising during execution ^7^.

At any trial t, participants plan to exert a force (motor command) *u*_*t*_. Motor execution noise 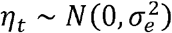 corrupts the movement, which ends up at a position *x*_*t*_ :

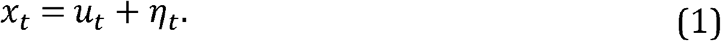

The movement error *δ*_*t*_ is defined as the difference between the target position *z*_*t*_ and the actual marble endpoint x_t_. The perceived error 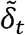 is, however, also corrupted by the sensory noise 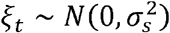 :

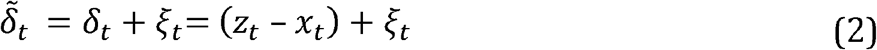

Finally, this subjective error is used to correct the motor plan at an adaptation rate *α* :

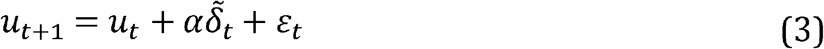

Where 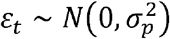 is a noise potentially corrupting motor plans between trials. Given the execution and sensory noise variances, we define execution and sensory precision as their respective inverses, 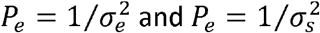.

Note that replacing Eqns. (1) and (2) in Eqn. (3) reveals that larger adaptation rates amplify the influence of execution and sensory noise and that sensory noise induces a stochastic drift in force production.

Because the model is linear-Gaussian, its predictive distribution is fully characterised by its mean and variance, which we derive analytically below.

As the goal of the task is to reach the target (δ =0), we also identified the optimal learning rate which minimises the expected error in the long run (Kalman gain):

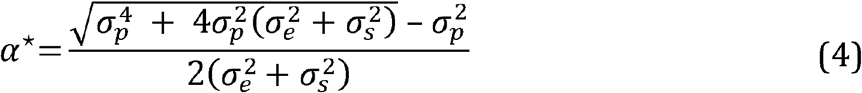

Notably, the optimal adaptation rate decreases with both execution 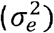 and sensory 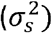 noise. For each fitted participant and condition, we calculated the corresponding normative adaptation rate from the estimated execution and sensory noise parameters. State noise, which was not estimated from the data, was arbitrarily set to 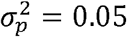.

#### Derivation of the static model

To make our computational model more tractable, we derive the mean of the predicted motor performance as a function of trial, yielding a deterministic prediction function that can then be fitted to the empirical performances.

To this end, let’s first rewrite the evolution function (Eqn. 3) by substituting *x* and 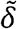 from the observation and error functions (Eqns. 1 and 2):

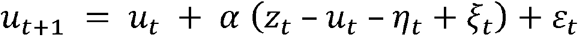

To obtain a deterministic description of the stochastic dynamics, we derived the mean and variance of the motor plan *u*_*t*_. Denoting *E[u*_*t*_*] = m*_*t*_, we have:

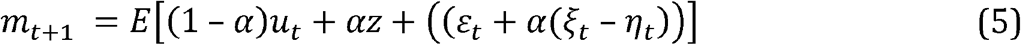

which yields, by recursion:

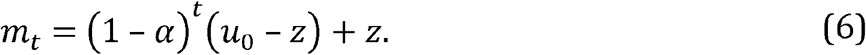

Now denoting *Var(u*_*t*_*) = v*_*t*_, we have:

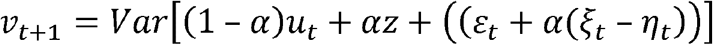

Solving the recursion gives:

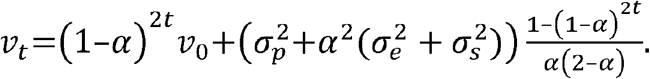

The predictive distribution of the motor plan is therefore characterised by its mean and variance at each trial.

In our behavioural task, we cannot directly assess the motor command *u*_*t*_ and can only measure the actual performance *x*_*t*_ and assess how far it falls from the target, as captured by *δ*. Plugging Eqn. 1 into 2 gives:

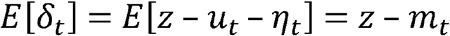

and

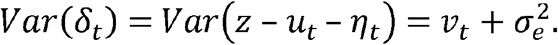

Using 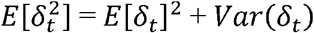 and *ν*_o_ = 0, we can now write the expected distance to the target for a given trial *t*, which can then be fitted to empirical data

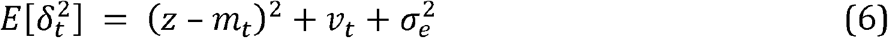

<h3>Limit behaviour

Note that as the number of trials *t* increases, the average performance *m*_*t*_ will converge to the target:

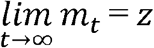

and the variance will reach a steady state:

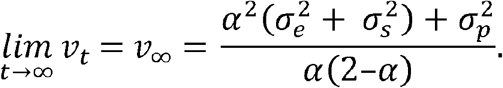

Replacing the above in Eqn. 7 relates to the expected distance to the target after learning:

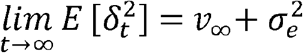

Setting the derivative to zero provides the optimal learning rate that minimises the asymptotic expected error given in Eqn. 4.

### Model fitting

Model prediction of normalised aiming errors 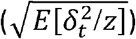 was fitted to empirical error trajectories separately for each bloc (30 trials) of the 8 possible conditions (target position × valence × feedback blinding) using variational Bayesian inference as implemented in the VBA toolbox ^64^. Fitting was done in log space to temper the influence of initial trials. Because motor noise is known to scale linearly with force, we reparametrized the model using 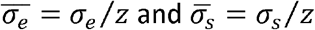 to derive force-invariant noise estimates (noise scaling parameters). State noise 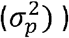 was arbitrarily fixed to 0 to ensure identifiability. With the adaptation rate a, the model therefore included three free parameters. The sensory noise 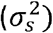 was set to zero in blocks with visual feedback.

We first estimated group-level parameters by fitting the squared error time series averaged across all participants within each group. In order to reduce bias, we then derived individual parameter estimates for each participant i using Jackknife resampling:

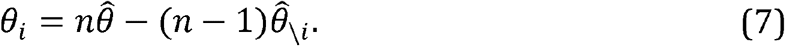

where θ is any free parameter of the model,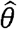 and 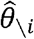 are the parameter estimates using the average behaviour across all participants of the group with and without the participant *i*, respectively, and is the number of participants in the group.

All model-based analyses were performed using MATLAB R2023a (The MathWorks Inc., Natick, MA).

## Data availability

The human participant-level data reported in this study cannot be deposited in a public repository under the European Union General Data Protection Regulation and the Institutional Review Board data protection policies. To request access, please contact the lead contact. Data provision may include processed and unprocessed data and will require a data-sharing agreement. Data sharing requires that the purpose of data reanalysis align with the study aims, as approved by the ethics review boards and participants. Furthermore, consent to data privacy must be obtained by signing the agreement form. Requests will be answered within 4 weeks. Source data for generating the figures are provided with this paper.

## Acknowledgment

We thank Y. Saadi and E. Alberti for assisting with the data acquisition. Funding: The study was supported by the Deutsche Forschungsgemeinschaft (DFG; German Research Foundation) under Germany’s Excellence Strategy – EXC 2030– 390661388 and through SFB 1451, project ID 431549029, C06 (F.J. and M.T.) and B05 (L.V.), as well as by the German Centre for Diabetes Research, project ID 82DZD05H1G (M.T.).

## Author Contributions

L.R., F.J., and M.T. designed experiments. L.R. designed the task and constructed the modelling analysis together with P.E.Z., who programmed the analysis chain. P.Z. recruited the patients, performed the clinical assessments, and supported the experiments as study physicians. P.E.Z. and L.R. analysed the data. L.R., M.T., T.B., and L.J.V. interpreted the data. L.R. and M.T. wrote the manuscript with input from all other authors.

## Competing Interests

The authors declare no competing interests.

## Figure legends

**Extended Data Figure 1.**
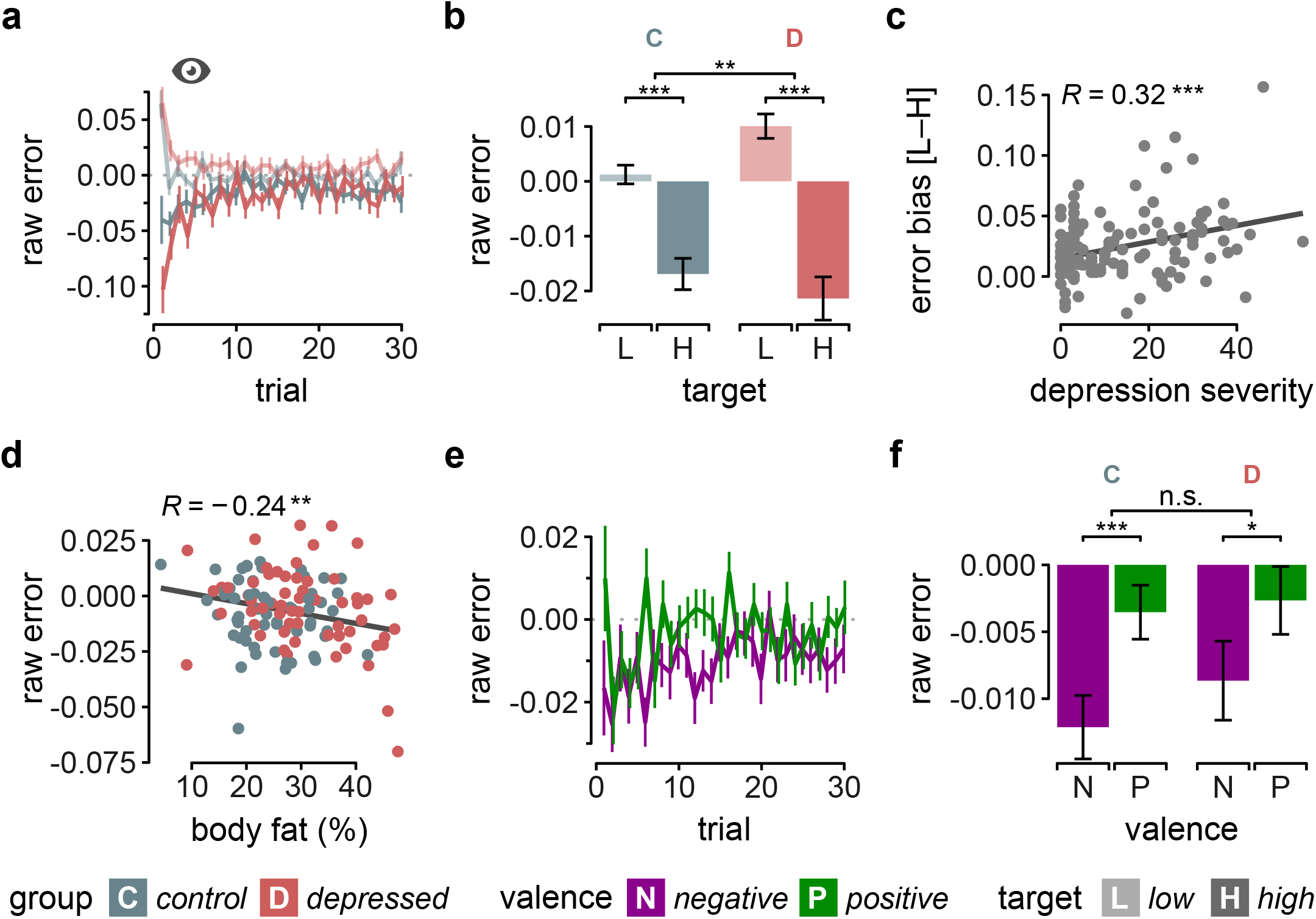
Raw aiming errors across force and valence conditions. a, Dynamics of raw (signed) aiming error under visual feedback for low- and high-target conditions in healthy controls and patients with depression. b, Mean raw aiming error under visual feedback for low- and high-target conditions in healthy controls and patients with depression. c, Relationship between depression severity (BDI-II score) and the difference in raw aiming error between high- and low-target conditions. d, Relationship between body-fat percentage and raw aiming error. e, Dynamics of raw aiming error under visual feedback for positive- and negative-valence conditions. f, Mean raw aiming error under visual feedback for positive- and negative-valence conditions in healthy controls and patients with depression. Negative-valence (N) and positive-valence (P) conditions are shown in purple and green; low-target (L) and high-target (H) conditions are shown in lighter and darker shades; healthy controls (C) and patients with depression (D) are shown in blue and red. Data are means ± SEM across participants. Relationships in c,d are quantified using Pearson correlations with maximum-likelihood regression lines. Statistical comparisons were performed using linear mixed-effects models with ANOVA for main effects and interactions and paired or unpaired t-tests for pairwise comparisons, as appropriate. Significance levels: p* < .05, p** < .01, p*** < .001.

**Extended Data Figure 2.**
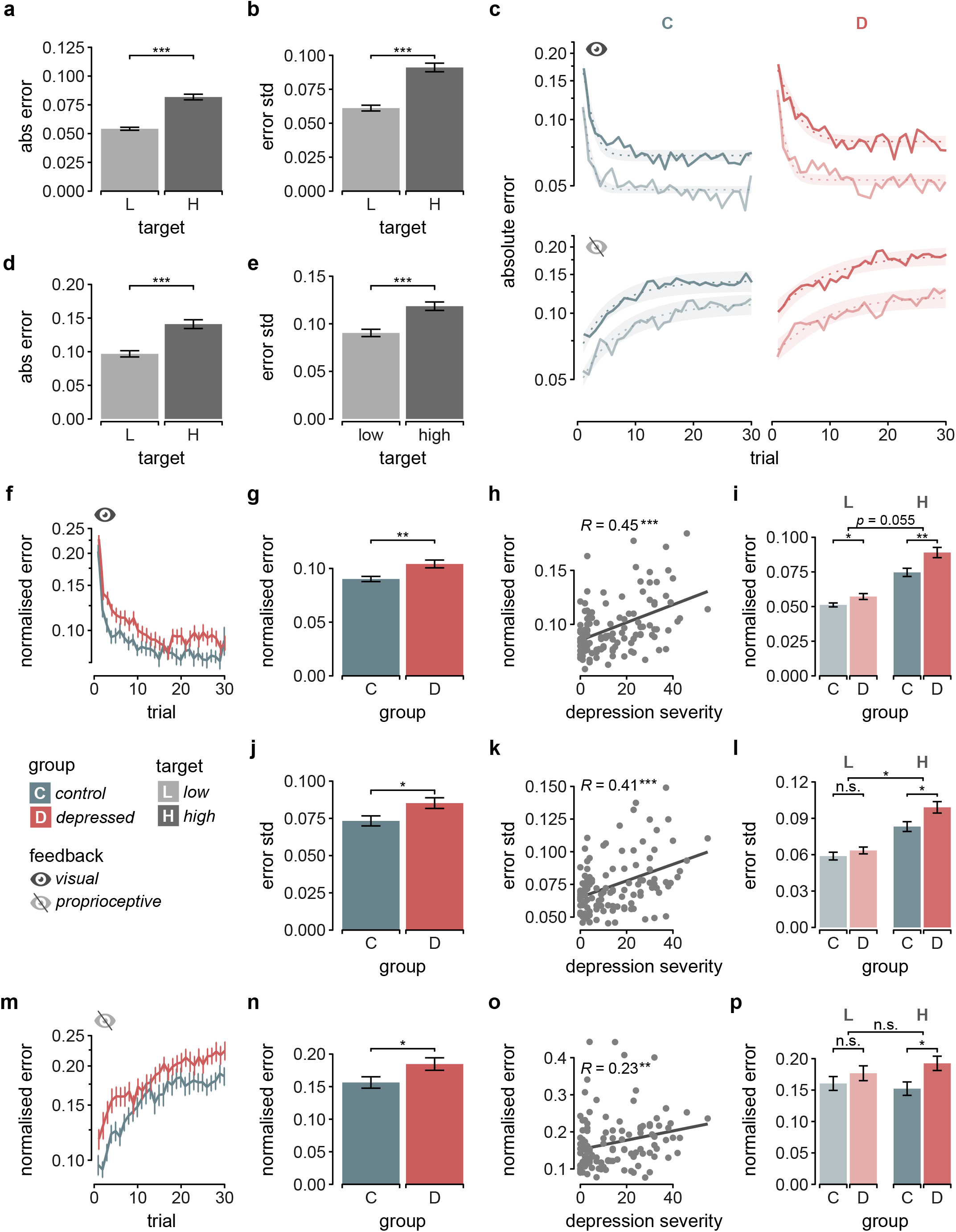
Additional analyses of aiming error. a, Mean absolute aiming error under visual feedback for low- and high-target conditions. b, Standard deviation of aiming error under visual feedback for low- and high-target conditions, averaged across trials. c, Model fits to absolute aiming-error trajectories. Left and right columns show healthy controls and patients with depression, respectively; upper and lower rows show adaptation with and without visual feedback, respectively. Solid lines are empirical data. Dotted lines and shaded area represent the mean and 95% confidence interval of the (posterior) model predictions. d, Mean absolute aiming error without visual feedback for low- and high-target conditions. e, Standard deviation of aiming error without visual feedback for low- and high-target conditions. f, Adaptation trajectories of normalised aiming error under visual feedback for healthy controls and patients with depression. Normalised aiming error was calculated as the square root of squared aiming error divided by target amplitude. g, Mean normalised aiming error under visual feedback for healthy controls and patients with depression. h, Relationship between depression severity (BDI-II score) and normalised aiming error under visual feedback. i, Mean normalised aiming error under visual feedback for healthy controls and patients with depression, shown separately for low- and high-target conditions. j, Standard deviation of aiming error under visual feedback for healthy controls and patients with depression. k, Relationship between depression severity (BDI-II score) and aiming-error variability under visual feedback. l. Standard deviation of aiming error under visual feedback for healthy controls and patients with depression, shown separately for low- and high-target conditions. m, Normalised aiming-error drift without visual feedback for healthy controls and patients with depression. n, Mean normalised aiming error without visual feedback for healthy controls and patients with depression, averaged across trials. o, Relationship between depression severity (BDI-II score) and normalised aiming error without visual feedback. p, Mean normalised aiming error without visual feedback for healthy controls and patients with depression, shown separately for low- and high-target conditions and averaged across trials. Low-target (L) and high-target (H) conditions are shown in lighter and darker shades; healthy controls (C) and patients with depression (D) are shown in blue and red. Standard deviation in b, e, j-l is computed across the last 15 trials of each block (asymptotic behaviour) and averaged across conditions. Data are means ± SEM across participants. Relationships in h, k and o are quantified using Pearson correlations with maximum-likelihood regression lines. Statistical comparisons were performed using linear mixed-effects models with ANOVA for main effects and interactions and paired or unpaired t-tests for pairwise comparisons, as appropriate. Statistical tests and p-values are reported in the main text or Methods. Significance levels: p* < .05, p** < .01, p*** < .001.

**Extended Data Figure 3.**
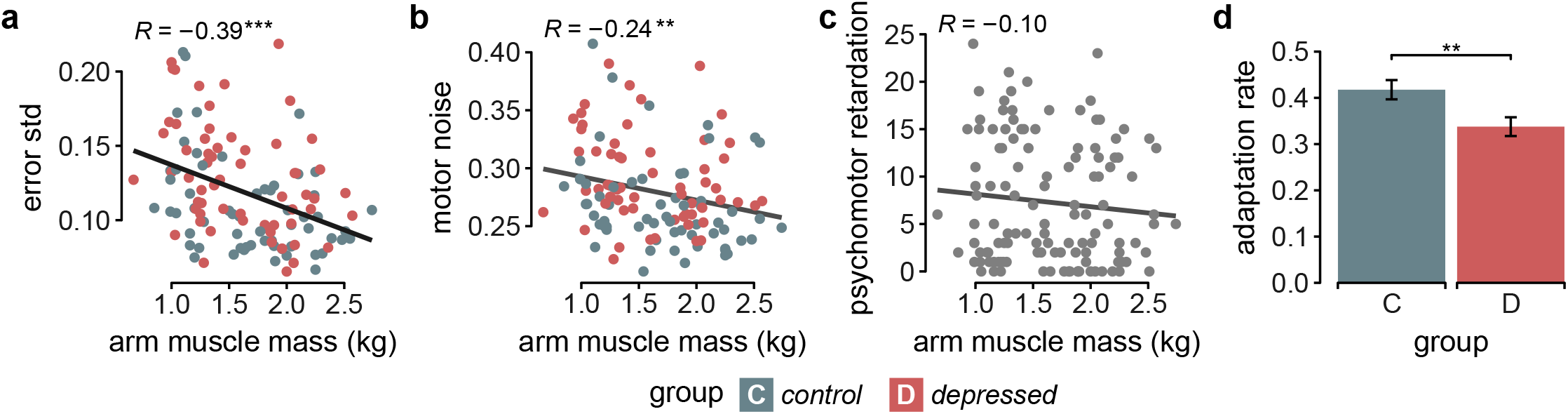
Muscle mass relates to motor variability but not psychomotor retardation. a, Relationship between aiming-error variability (i.e. standard deviation of absolute errors computed across all trials and averaged across conditions) and muscle mass of the arm used for the task, measured by bioelectrical impedance analysis. b, Relationship between estimated motor execution noise and muscle mass of the task-performing arm. c, Relationship between psychomotor retardation (SRRS score) and muscle mass of the task-performing arm. d, Adaptation rate in healthy controls and patients with depression. Muscle mass is reported in kg. Healthy controls (C) and patients with depression (D) are shown in blue and red. Relationships in a–c are quantified using Pearson correlations with maximum-likelihood regression lines. Data in d are means ± SEM and compared using a t-test. Statistical tests and p-values are reported in the main text or Methods. Significance levels: p** < .01, p*** < .001.

**Extended Data Figure 4.**
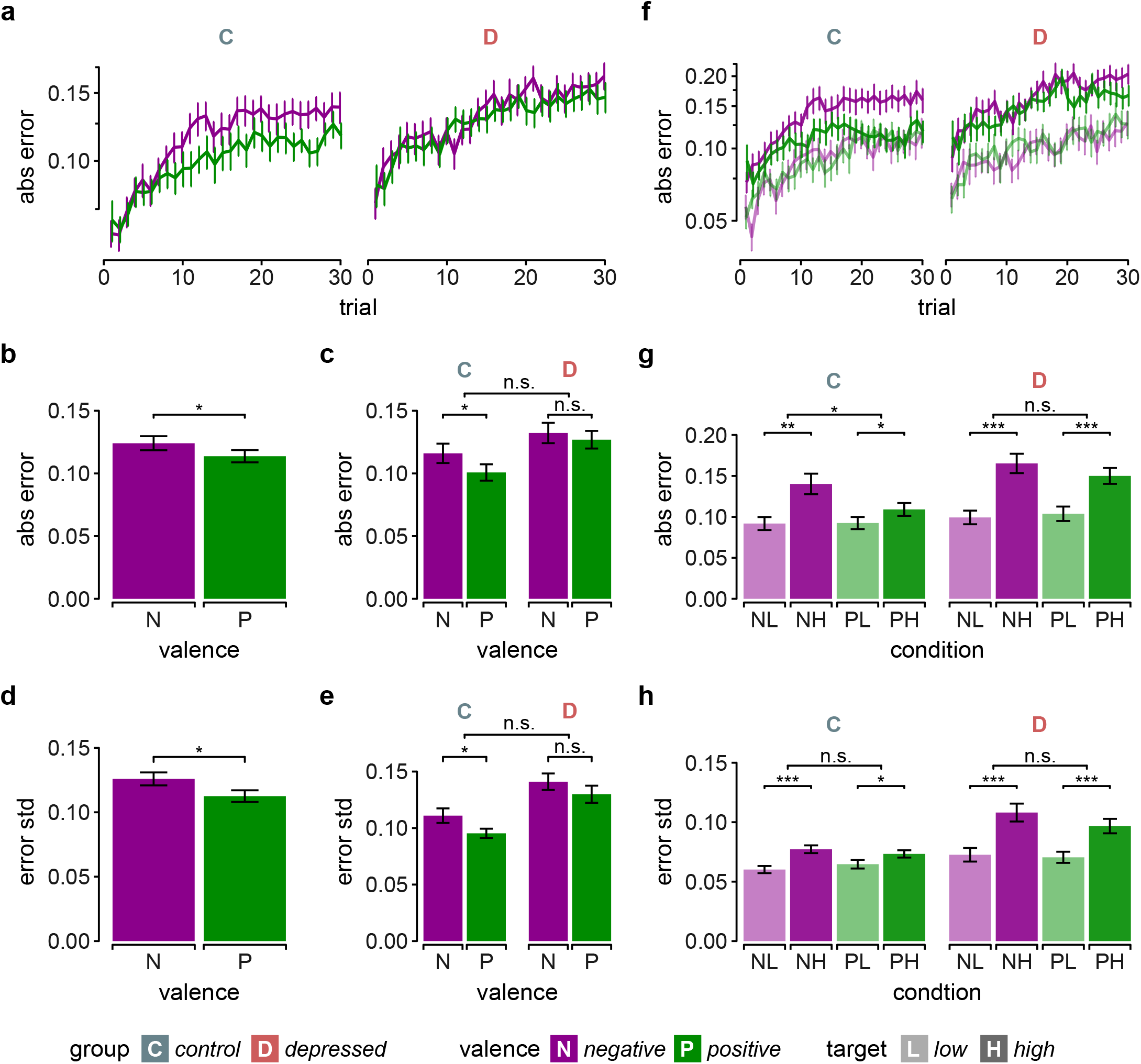
Effects of feedback valence on proprioceptive performance. a, Aiming-error trajectories without visual feedback for positive- and negative-valence conditions in healthy controls and patients with depression. b, Mean absolute aiming error without visual feedback for positive- and negative-valence conditions. c, Mean absolute aiming error without visual feedback for positive- and negative-valence conditions in healthy controls and patients with depression. d, Standard deviation of aiming error without visual feedback for positive- and negative-valence conditions. e, Standard deviation of aiming error without visual feedback for positive- and negative-valence conditions in healthy controls and patients with depression. f, Aiming-error trajectories without visual feedback for positive- and negative-valence conditions, shown separately for low- and high-target conditions in healthy controls and patients with depression. g, Mean absolute aiming error without visual feedback for positive- and negative-valence conditions, shown separately for low- and high-target conditions in healthy controls and patients with depression. h, Standard deviation of aiming error without visual feedback for positive- and negative-valence conditions, shown separately for low- and high-target conditions in healthy controls and patients with depression. Negative (N) and positive (P) valence are shown in purple and green. Low (L) and high (H) targets are shown in lighter and darker shading. Healthy controls (C) and patients with depression (D) are shown in blue and red. Standard deviation in d,e,h is computed across the last 15 trials of each block (asymptotic behaviour) and averaged across conditions. Data are means ± SEM. Statistical comparisons were performed using linear mixed-effects models with ANOVA for main effects and interactions and paired or unpaired t-tests for pairwise comparisons, as appropriate. Significance levels: p*< .05, p** < .01, p*** < .001.

## Notes

### Competing Interest Statement

The authors have declared no competing interest.

