## Supplementary Table 1 for "Motor learning adapts to effort-dependent uncertainty"

|  |  | **clinical scale** | | | |
| --- | --- | --- | --- | --- | --- |
| **behavioural marker** | **figure** | BDI-II | MADRS | DARS | SRRS |
| execution noise | 1i | 0.37*** | 0.34*** | –0.28** | 0.26** |
| sensory noise [H-L] | 1l | 0.36*** | 0.28** | –0.34*** | 0.23*** |
| optimal adaptation rate | 2h | –0.42*** | –0.39*** | 0.32*** | –0.32*** |
| adaptation rate interaction | 3h | –0.24** | –0.30** | 0.17° | –0.19* |
| skeletal muscle mass | ED 3c | –0.17° | –0.16 | 0.07 | –0.09 |

***Correlation between behavioural markers and clinical scales****. ***: p* < .001*, **: p* < .01*, *: p* < .05*,* °*: p* < .1*,*
